# Development of a mouse model to investigate the effect of exertional heat stress on skeletal muscle

**DOI:** 10.64898/2026.10.27.762193

**Authors:** Yifan Chen, Richard C. Steiner, Tianzheng Yu

**Author notes:** Corresponding author: Yifan Chen, Ph.D., Department of Military and Emergency Medicine, Uniformed Services University, 4301 Jones Bridge Road, Bethesda, MD 20814, USA.

## Abstract

Increased body temperature during exercise is considered a risk factor for skeletal muscle injury. There are various murine models of heat exposure-induced organ injury. The present study has attempted to develop a mouse model of skeletal muscle injury associated with exercise-induced whole body hyperthermia. C57BL/6J male mice were randomly assigned into control (CON) and exercise (EX) groups (n = 6 per group). EX group exercised on a programmed, motorized wheel, while CON group remained in a mouse cage in an environmental chamber at 35°C for ~3 hours. EX group showed a significantly higher average body core temperature (38.5°C ± 0.3) compared to CON group (37.3°C ± 0.3). Plasma myoglobin levels were significantly higher in EX group than in CON. Additionally, exercise significantly impaired skeletal muscle and mitochondria, as evidenced by elevated TBARS and caspase-3 levels and reduced ATP and Complex IV levels in gastrocnemius muscles in EX group. We developed a mouse model of exercise-induced mild whole body hyperthermia that recapitulates many key features of heat stress-induced skeletal muscle injury. This novel model will provide a useful tool for future investigation into the mechanisms of and intervention strategies for muscle exertional heat injury.

## 1. Introduction

Prolonged and strenuous physical exercise can cause skeletal muscle damage (1,2), or even rhabdomyolysis (massive amounts of muscle fibers breakdown) particularly in hot climates (3), which is a serious threat to military readiness (4). Increased body temperature (exertional heat stress) is considered a contributor to exercise-related muscle injury (3). Our previous studies showed that heat exposure-induced hyperthermia is associated with skeletal muscle injury in mice (5,6). During exercise, body temperature may rise due to the muscle metabolic and contractile activities. Investigation into the mechanisms of and intervention strategies for muscle exertional heat injury is impeded by a lack of murine models.

In this study, we developed a mouse model that features mild exertional heat stress and key aspects of skeletal muscle injury. We confirmed that their elevated body temperature was a result of exercise by comparing with a sedentary control group in the same experimental condition. Then we examined the effects of exercise on muscle injury-related markers at the tissue and systemic levels.

## 2. Methods

### 2.1 Animals

The study protocol was approved by the Uniformed Services University (USU) Institutional Animal Care and Use Committee in accordance with all applicable Federal regulations governing the protection of animals in research (MEM-25-059). Male C57BL/6J mice (6 weeks of age) were purchased from Jackson Laboratories (Bar Harbor, ME) and were on chow diet in a controlled (21 °C) and enriched environment at the USU Department of Laboratory Animal Resources with a 12/12-hour light-dark cycle. All mice were surgically implanted with a temperature-sensing passive integrated transponder (PIT), Biotherm 13 PIT Tag (Biomark Inc, Boise, ID) in the abdominal cavity under anesthesia. At least two weeks were allowed for recovery. Mice were randomly assigned to control (CON) and exercise (EX) groups (6 per group).

### 2.2 Mouse running wheel controlling and body temperature recording

As shown in Fig. 1, the system comprises a custom-designed brushless motor-driven wheel and a PIT tag reader (Biomark, Inc., Boise, ID). The wheel is made of plexiglass to allow transmitting signals to the reader from the mouse tag. The motor is connected to a DC motor driver. A custom-developed LabView (National Instruments, Austin, TX) interface was used to control the motor driver and record the motor speed and tag data in a Windows-based PC.

**Fig. 1.**
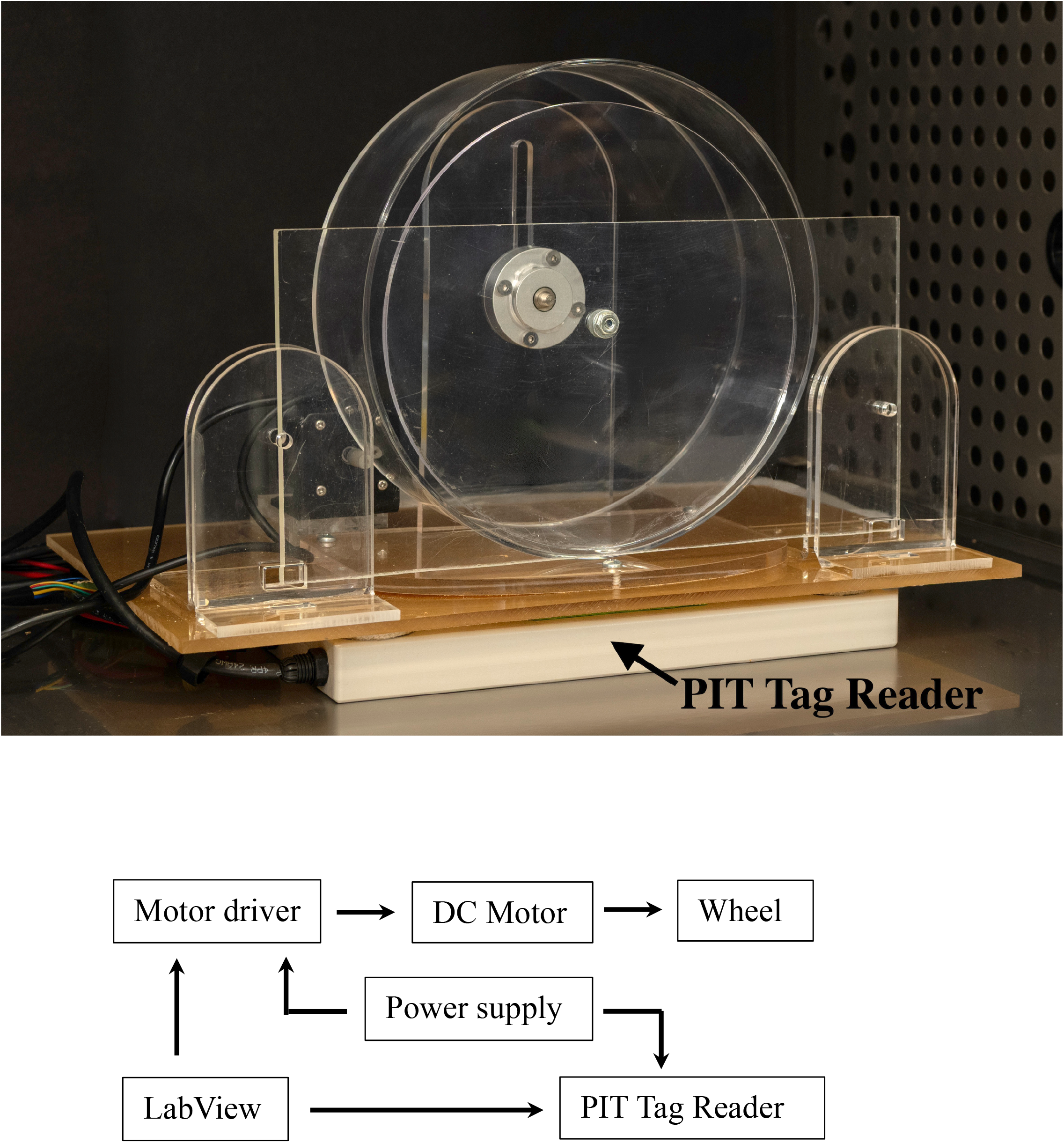
Custom-designed motorized-wheel system. The main components include a modified acrylic plexiglass mouse exercise wheel, a brushless motor, a DC motor driver, and a PIT tag reader. The motor and reader are powered by a DC source. A custom-developed LabView interface was used to control and record the motor speed and PIT tag data in a Windows-based PC.

### 2.3 Wheel running

#### Training

Both CON and EX mice underwent three training sessions to familiarize to the mouse running wheel, with an interval of 2 - 3 days. Each session included 10 minutes of voluntary wheel running (motor disengaged) and 30 minutes of forced wheel running at a speed of 2 - 4 m/min. The final session was performed in an environmental chamber (Model 3950, Thermo Forma, Marietta, OH) at room temperature, followed by three days of wash-out before experiments.

#### Experiments

For each experiment, 1 CON and 1 EX mice were placed in a mouse cage with food and water in the chamber at room temperature overnight, and their body core temperature (Tc) and activity were continuously recorded. The experiment was performed on the following morning from ~0830 to1200. Food and water were removed before turning on the chamber heating mechanism. In our preliminary study, we exposed mice to two chamber temperatures, 36.5 °C and 35.0 °C, for 3 hours. The results showed that the Tc increased (peak Tc: 39.0 - 39.4°C; average Tc: 38.4 - 38.5°C) at 36.5 °C, but not at 35.0 °C. Thus, the experimental chamber temperature was preset at 35.0 °C. After the chamber temperature was reached, the EX mouse was transferred to the wheel.

Running wheel speed began at 2.2 m/min with increments of 0.2 m/min every 5 min until reaching 6 m/min and would continue for up to 200 min. The CON mouse remained in the cage alongside the EX mouse in the 35.0 °C chamber for the entire exercise period. Immediately after the experiments, blood was collected by cardiac puncture into a heparin-coated tube under anesthesia and plasma was separated by centrifugation. Gastrocnemius muscles were removed after euthanasia. All samples were flash-frozen in liquid nitrogen, and stored at −80 °C.

### 2.4 Plasma analysis

Myoglobin and creatine kinase MM were measured using the Mouse Myoglobin ELISA Kit (CrystalChem, CAT #80654, Itasca, IL), and Mouse Creatine Kinase ELISA Kit (Invitrogen, CAT #EEL217, Waltham, MA), respectively, according to the manufacturers’ instructions.

### 2.5 Gastrocnemius muscle analysis

Tissue samples were homogenized in respective assay buffer on ice, centrifuged at 4°C, and the supernatants were collected. The concentrations of thiobarbituric acid reactive substances (TBARS), activated caspase-3, adenosine triphosphate (ATP), and mitochondrial complex IV (cytochrome c oxidase) enzyme were measured in gastrocnemius muscle tissue using the following assay kits, per the manufacturers’ instructions: TBARS (Cayman Chemical, CAT #10009055, Ann Arbor, MI), Human/Mouse Cleaved Caspase-3 (Novus Biologicals, CAT #DYC835-2, Centennial, CO), ATP (Abcam, CAT #ab83355, Cambridge, United Kingdom), and Cytochrome Oxidase Activity Colorimetric (BioVision, CAT #K287, Exton, PA). For all assays, samples were run in duplicate, absorbance was measured using the iMark™ Microplate Absorbance Reader, (Bio Rad, Hercules, CA), and values were calculated from standard curves and normalized to total protein concentration.

### 2.6 Statistical analysis

All data are expressed as the mean ± SD and analyzed using GraphPad Prism 10.4. Significance tests were two-tailed unpaired t and two-way ANOVA with Tukey’s post-hoc. In all statistical comparisons, a P-value < 0.05 was considered significant.

## 3. Results

### 3.1 Exercise induces an increase in Tc

No differences in average Tc were found between EX and CON groups (EX: 37.4°C ± 0.5 vs CON: 37.3°C ± 0.5, P > 0.05) overnight before their experiments at the room temperature 21°C (Fig. 2). At the experimental temperature 35°C, EX and CON groups had an average Tc of 38.5 ± 0.4 °C (peak Tc: 39.3°C ± 0.4 °C) and 37.3°C ± 0.3 °C during up to 200 min wheel exercise or sedentariness, respectively. These results indicate that exercise induced mild whole body hyperthermia in mice.

**Fig. 2.**
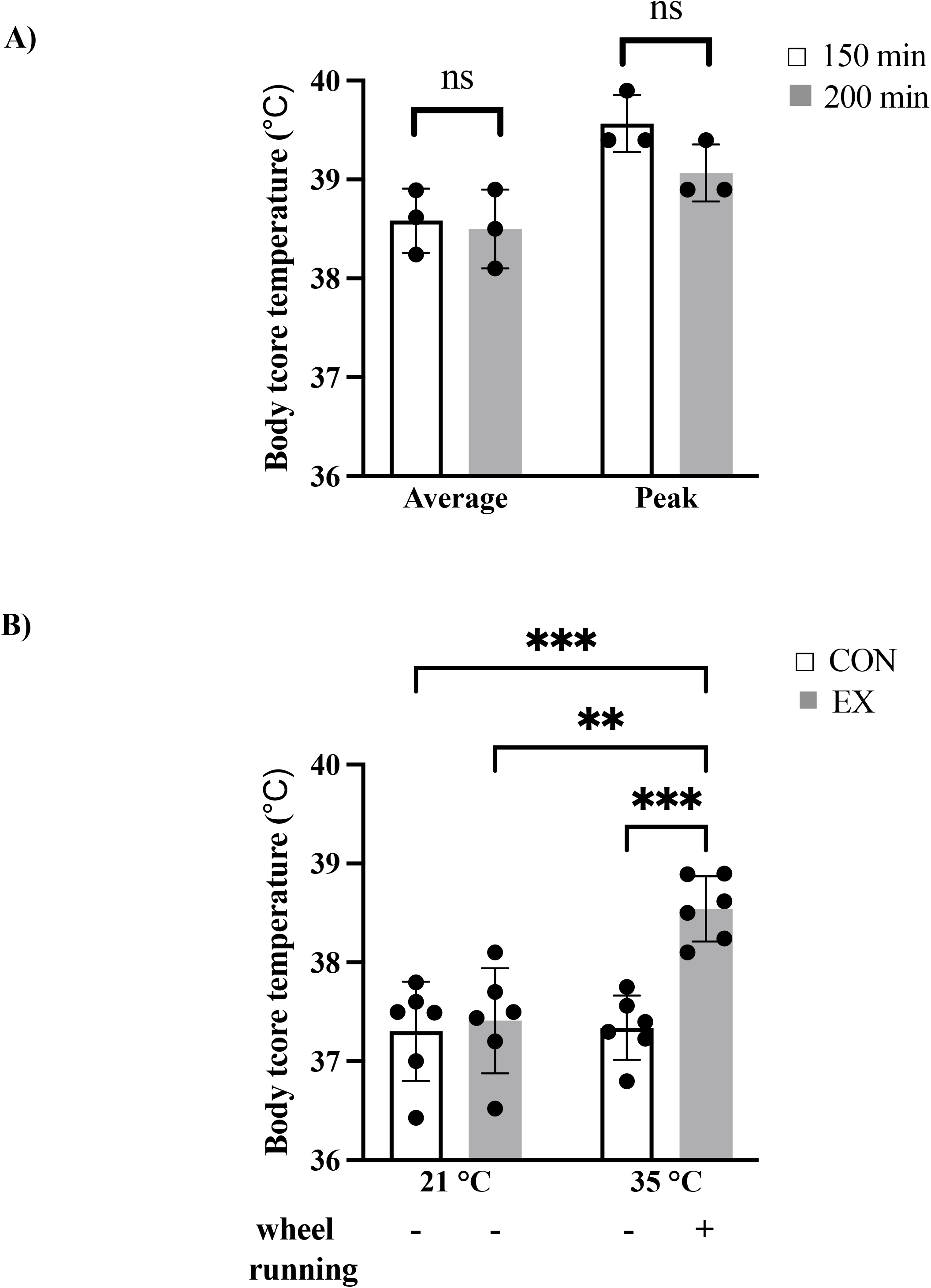
Average body core temperature (Tc) in CON and EX groups at 21°C and 35°C. ** P < 0.01 and *** P < 0.001 by two-way ANOVA. Data are shown as mean ± SD (n = 6 per group).

### 3.2 Exercise increases circulating muscle injury marker levels

Skeletal muscle expresses myoglobin and two isoenzymes of creatine kinase (CK), CK-MM (dominant) and CK-MB. Both myoglobin and CK are cytoplasmic proteins and can enter the systemic circulation upon cell damage. Circulating myoglobin rises within hours, while CK enters the bloodstream with a delay due to lymphatic transport, peaks 2–4 days after muscle tissue injury (7). In this study, immediately after exercise, EX group showed a significantly increased level of plasma myoglobin compared to CON group; whereas no differences in plasma CK-MM were detected between the two groups (Fig. 3).

**Fig. 3.**
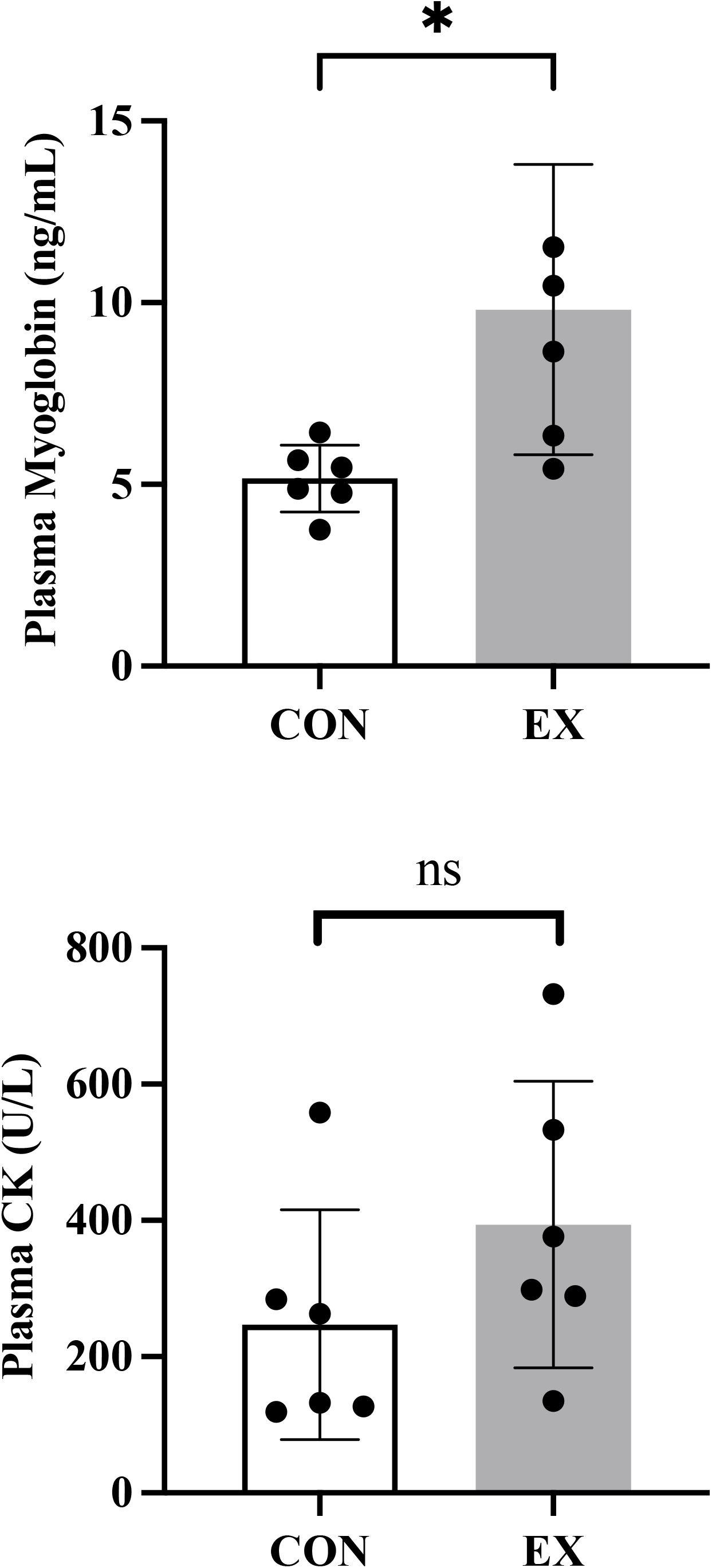
Plasma myoglobin and creatin kinase concentrations in CON and EX groups. Data are shown as mean ± SD (n = 6 per group). * P < 0.05 by a two-tailed unpaired t test. NS: no significance.

### 3.3 Exercise causes muscle oxidative stress, apoptosis, and mitochondrial impairment

Tissue TBARS measures oxidative stress-induced lipid peroxidation, while caspase-3 acts as a central mediator of apoptosis. TBARS and caspase-3 are key biomarkers of cellular damage. As shown in Fig. 4, EX group showed significantly increased levels of TBARS and caspase-3 in gastrocnemius muscles compared to CON group, indicating that exercise caused muscle damage.

**Fig. 4.**
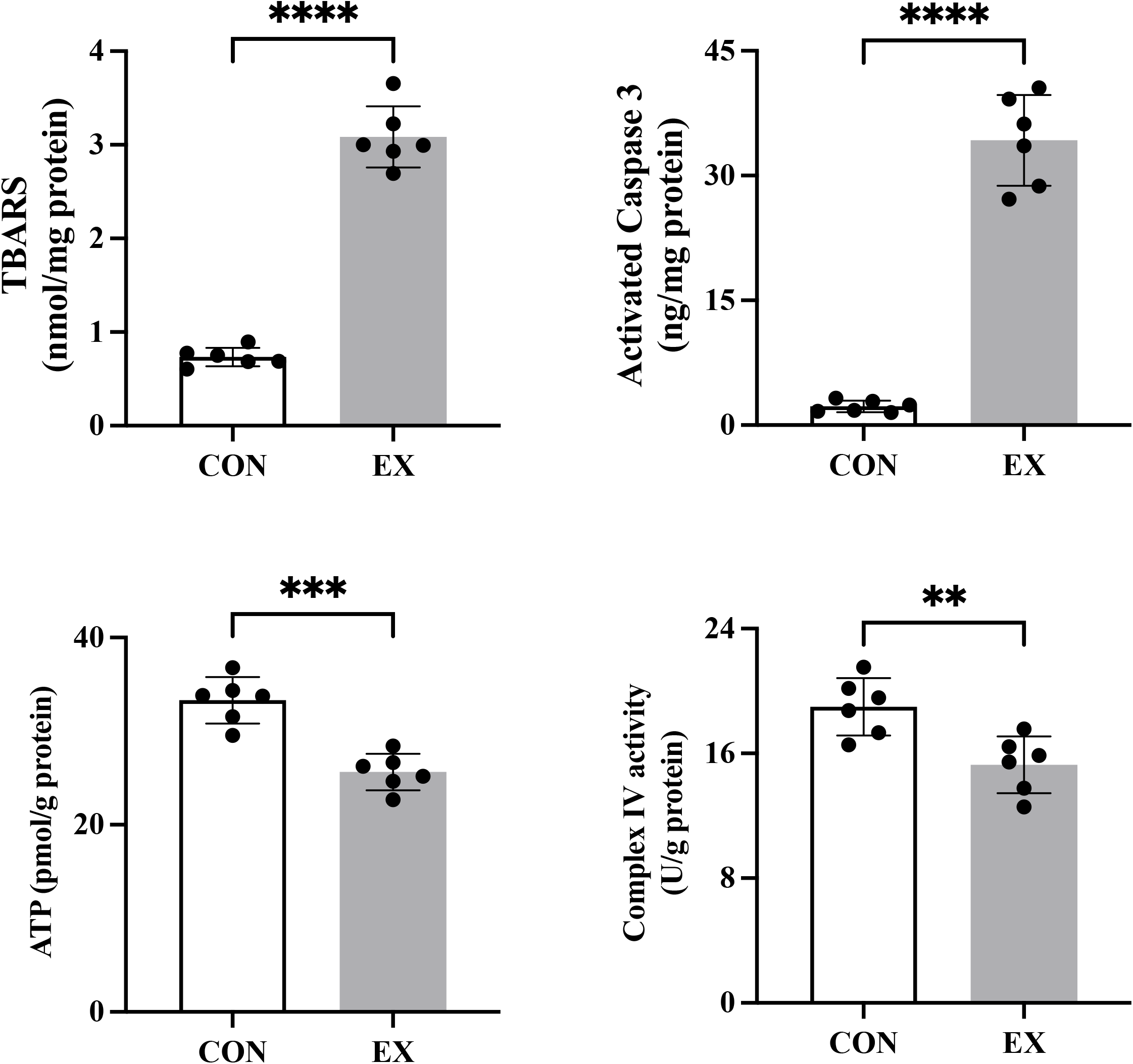
Gastrocnemius muscle TBARS, capsase-3, ATP, and Complex IV levels in CON and EX groups. Data are shown as mean ± SD (n = 6 per group). ** P < 0.01, *** P < 0.001, and **** P < 0.0005 by a two-tailed unpaired t test.

Tissue ATP and Complex IV (cytochrome c oxidase) levels are tightly coupled in mitochondrial oxidative phosphorylation. They serve as the key determinants of cellular energy status. Following exercise, EX group had significantly reduced levels of ATP and Complex IV in gastrocnemius muscles compared to CON group. These results indicate that exercise caused muscle mitochondrial impairment or energy depletion.

## 4. Discussion

In this study, we established a mouse model of exercise-induced mild whole body hyperthermia that features skeletal muscle injury. We demonstrated that increased body temperature in EX group was a result of exercise by comparing with the sedentary CON group exposed to the same experimental condition. A number of mouse models of exertional heat stroke have been created resulting from exercise at a temperature of 37.5 - 39.5 °C (8-10). To our knowledge, this study is the first to reveal the presence of hyperthermia and skeletal muscle injury associated with exercise at an ambient temperature below the normal physiological range in mice.

The principal finding of this study is that a single bout of dynamic exercise induced skeletal muscle impairment, as evidenced by increased levels of circulating myoglobin and muscle oxidative stress and apoptotic markers, and reduced levels of muscle ATP and Complex IV. Our exercise protocol is considered low intensity with a long duration, whereas high intensity short duration exercise may produce a detrimental effect in mice (11). As a relevant limitation, we did not conduct a follow-up study. Thus, we do not know whether the exercise-induced alterations in this study are reversible.

Exercise-related muscle injury, including rhabdomyolysis, is likely a result of combined mechanical (contractile) and thermal effects on the working muscles, although the underlying mechanisms are not fully understood. The results of this study showed that exercise-induced muscle apoptosis was associated with mitochondrial impairment and oxidative stress. Mitochondria play a central role in stress-induced reactive oxygen species production and apoptosis (12). Increased temperature has been shown to impair mitochondrial function or efficiency in skeletal muscle in mice (5), rats (13), dogs (14), and humans (15). Our findings of reduced muscle ATP and Complex IV content in EX mice indicate that exercise caused muscle mitochondrial impairment. As a result of reduced efficiency of ATP synthesis, increased energy demand or heat production through oxidative metabolism is needed to maintain the desired level of ATP synthesis. Skeletal muscle overheating can lead to increased production of reactive oxygen species that causes oxidative cell membrane damage (increased TBARS). Together, our results are consistent with oxidative stress-related energy metabolic alterations, suggesting the association between mitochondrial impairment and exercise-induced skeletal muscle injury.

There are several limitations of this study. First, we were unable to separate thermal effects from exercise effects. It is difficult to examine these two effects independently, given that skeletal muscle activity is inherently associated with increased skeletal muscle temperature due to the metabolic activity needed to support skeletal muscle contraction. We do not know whether mice could complete the same exercise protocol at a much lower ambient temperature without increasing Tc; it would help determine the effects of exercise by comparing two groups that complete the exercise at the same intensity and duration, one with versus one without a hyperthermic response. Second, we only measured the temperature in the body, but not in the working muscles. It is very likely that the temperature increase was greater in working muscles than that (mostly due to muscle thermogenesis) in the whole body of EX mice during exercise. Unfortunately, measuring muscle temperature in free moving mice technically remains difficult. It is also interesting to note that the mildly elevated body temperature in EX group is similar to that reported in patients with rapid, severe muscle damage such as rhabdomyolysis (11). In addition, we observed that the Tc in EX group reached a plateau after ~150 min of exercise. These results suggest that exercise-induced muscle injury is associated with mild hyperthermia. Finally, considering the relatively small sample size and no female mice in this study, our findings may not be generalized. These limitations need to be addressed with further studies.

In summary, we developed a mouse model of exertional heat stress with key features of skeletal muscle injury that will facilitate the mechanistic study of the pathophysiology and development of intervention strategies of skeletal muscle exercise injury.

## Acknowledgements

This work was supported by the National Institutes of Health Office of Dietary Supplements (HU00012320098).

## Disclaimer statement

The opinions and assertions expressed herein are those of the author(s) and do not reflect the official policy or position of the Uniformed Services University or the Department of War. The contents of this publication are the sole responsibility of the author(s) and do not necessarily reflect the views, opinions or policies of The Henry M. Jackson Foundation for the Advancement of Military Medicine, Inc. Mention of trade names, commercial products, or organizations does not imply endorsement by the U.S. Government.

